# Temporal characteristics of gamma rhythm constrain properties of noise in an inhibition-stabilized network model

**DOI:** 10.1101/2022.11.23.517655

**Authors:** R Krishnakumaran, Supratim Ray

## Abstract

Gamma rhythm refers to oscillatory neural activity between 30-80 Hz, often induced in visual cortex by presentation of stimuli such as iso-luminant hues or gratings. Further, the power and peak frequency of gamma depend on the properties of the stimulus such as size and contrast. Gamma also has a typical arch shape, with a narrow trough and a broad peak, which can be replicated by a self-oscillating Wilson-Cowan (WC) model operating in an appropriate regime. However, oscillations in this model are infinitely long, unlike physiological gamma that occurs in short bursts. Further, unlike the model, gamma is faster after stimulus onset and slows down over time. Here, we first characterized gamma burst duration in LFP data recorded from two monkeys while they viewed full screen iso-luminant hues. We then added different types of noise in the inputs to the WC model and tested how that affected duration and temporal dynamics of gamma. While the model failed with the often-used Poisson noise, Ornstein-Uhlenbeck (OU) noise applied to both the excitatory and inhibitory populations in the WC model replicated the duration and slowing of gamma, and also replicated the shape and stimulus dependencies. Therefore, temporal dynamics of gamma oscillations put constraints on the type and properties of underlying neural noise.

## Introduction

Gamma rhythms refer to oscillatory activity in 30-80 Hz range generated by push-pull activity of excitatory and inhibitory neurons [1]. In the primary visual cortex (V1), gamma has been widely studied by presenting achromatic gratings and iso-luminant hue patches and has been shown to exhibit stimulus-dependent variations in power and frequency. For instance, larger stimuli produce gamma with higher power and lower frequency [2–4], whereas achromatic gratings presented at higher contrast with the same size generate gamma with higher peak frequency [5,6].

Gamma generation arising from push-pull activity has been demonstrated using different network models [7,8], with a recent study replicating the contrast response of gamma using a weak pyramidal interneuron network gamma (PING) model of V1 [9]. Another class of phenomenological models, called population rate models, lump excitatory and inhibitory neurons into a single population each (for example, the Wilson-Cowan (WC) models [10]). These models contain fewer free parameters and are more analytically tractable, and can exhibit a variety of network behaviors such as oscillations and hysteresis (see [10] for details). Recently, WC models of V1 have been demonstrated to explain size and contrast effects on gamma and population firing rates [11,12]. We have recently shown that the non-linear self-oscillating model described by Jadi and Sejnowski [11] (referred to here as the JS model) can also explain the non-sinusoidal ‘arch’ shape of the gamma rhythm in which the rhythm has a narrow trough and broad peak [13], but a stochastic WC model which is driven by Poisson inputs does not. Further, the JS model can also explain the strong reduction in gamma power in the presence of discontinuities in the stimulus [14]. However, gamma generated by the JS model is a limit cycle and therefore has infinite duration. On the other hand, experimental data shows that gamma oscillations due to achromatic gratings occur in bursts with a median duration of ~300 ms [15] or less [16] in V1, which is better explained by the stochastic noise-driven WC model [12]. Further, JS model produces gamma with a fixed frequency over the stimulus duration, while physiological gamma peak frequency is initially higher and decreases over time before reaching a plateau (for instance, see Fig 1D in [17], Fig 2B,E in [18], Fig 1 in [19], Fig 5 in [4], Fig 1 in [16] and Fig 1 in [20]).

**Figure 1:**
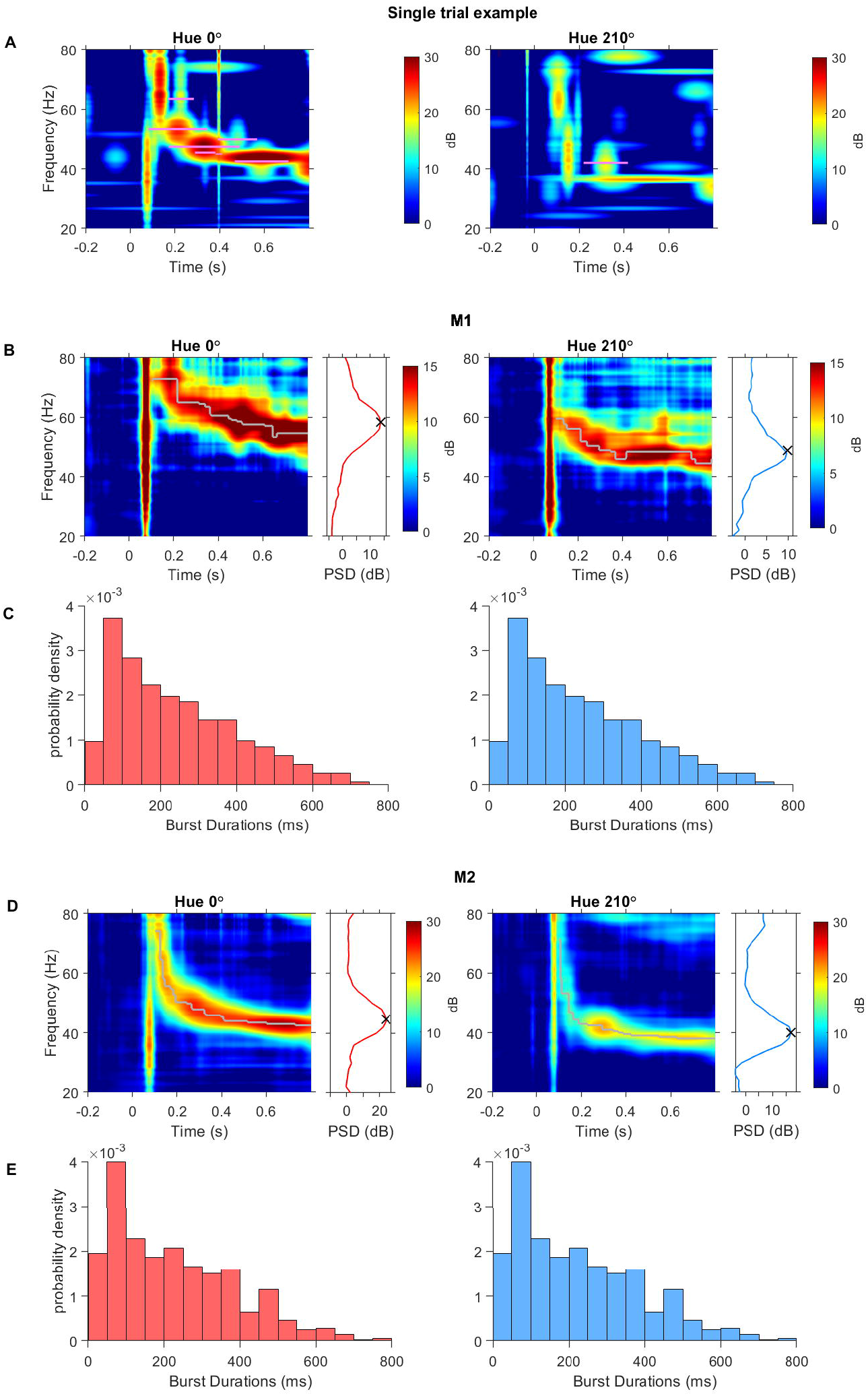
Gamma oscillations occur in bursts which exhibit a transient decrease in frequency after stimulus onset. (A) Single trial time-frequency spectrum corresponding to presentations of fullscreen red (left) and blue (right) hues. Horizontal magenta lines indicate the estimated location and duration of bursts. (B) Electrode and trial-averaged time-frequency spectrum from M1 corresponding to the example hues. Grey trace corresponds to the peak frequency of gamma at each timepoint after stimulus onset. The inset on the right of the time-frequency spectrum plots the corresponding electrode-averaged change in PSD (dB) between stimulus and baseline period with the detected gamma peak indicated by an ‘x’ mark. (C) Distribution of burst durations estimated in all trials and electrodes in M1. (D-E) Same as C-D for subject M2.

**Figure 2:**
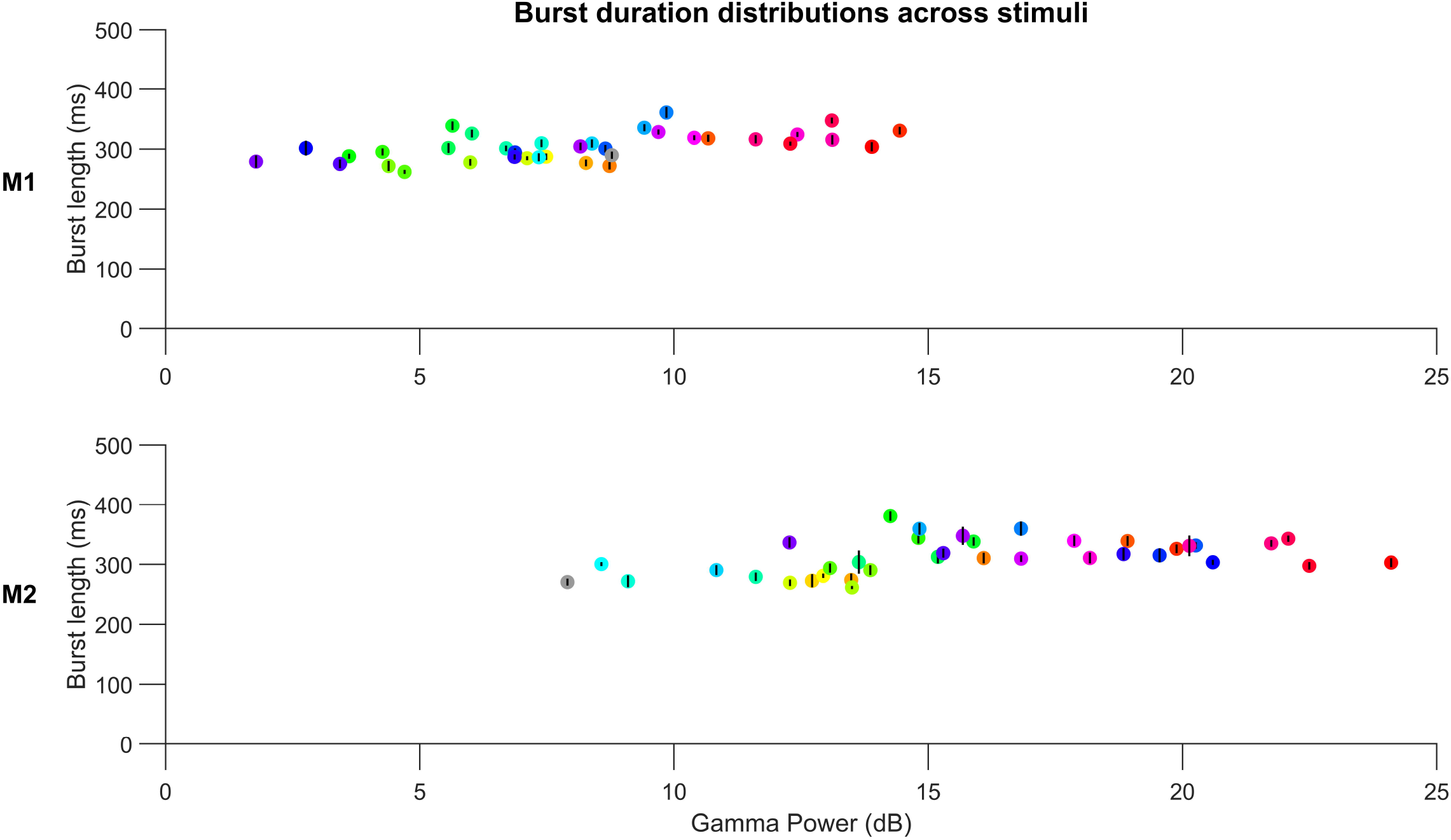
Median gamma burst durations are of the order of 300 ms across stimuli. Median of burst durations scattered against average gamma power for each stimulus from each monkey. Circles are colored to represent the presented hues while the grey circle represents the achromatic grating stimulus. Black error bars indicate the standard error of median (estimated by bootstrapping).

Bursty nature of gamma has inspired linear models in which the network acts like a damped oscillator perturbed by noisy inputs [16,17,21,22]. One such noise-driven piecewise linear model could demonstrate stimulus size and contrast dependence of gamma [12], although it could not exhibit the arch shape of gamma [13]. In contrast, noise driven non-linear models can produce bursty oscillations in the presence of noise either by damped oscillations set off by persistent noisy fluctuations or by noise-interrupted limit cycles [23]. Here, we tested whether addition of noise in the JS model could lead to generation of gamma oscillations with temporal characteristics observed in real data (appropriate burst duration and reduction of peak frequency after stimulus onset), in addition to maintaining the dependence on stimulus contrast and size as well as the arch shape of the rhythm, as demonstrated before.

## Results

### Burst duration analysis in LFP data due to full-screen iso-chromatic hue stimuli

We had previously shown that gamma bursts in V1 in response to full-screen achromatic grating stimuli have a duration of ~300 ms [15]. Here, we first computed gamma burst duration for full-field iso-luminant chromatic hues using the same approach. We presented full-screen patches of different hues to two monkeys, M1 and M2, and recorded LFP from 96 and 81 electrodes respectively. As in our previous study [13], we limited our analysis to 64 and 16 electrodes from the two monkeys which yielded reliable estimates of RF centers (see “Electrode selection” section in Materials and Methods).

Figure 1A shows the time-frequency spectrum of LFP recorded in an electrode during example trials where fullscreen red and blue hues were presented. Power in the gamma band occurred as bursts of different durations (the detected bursts are indicated by horizontal magenta lines, with length equal to the estimated duration of the bursts. For details, see *Time-Frequency analysis of LFP* in *Methods*). Figure 1B shows electrode and trial-averaged time-frequency spectra for the two hues from subject M1. The peak gamma frequency, indicated by the grey trace over the spectrum, was initially high and slowly decreased over a period of ~250 ms to a steady state value. This was the case for all the hues and the achromatic grating presentation. In the right subplots, the electrode and trial averaged PSD computed between 250-750 ms after stimulus onset is shown, exhibiting a distinct bump corresponding to the steady state gamma. Figure 1C shows the histogram of burst durations within the stimulus period for all trials and electrodes corresponding to the respective hue. Note that since the stimulus duration was only 800 ms and we only considered bursts which started and ended within the stimulus period, bursts longer than 800 ms were not considered. Nonetheless, the histogram exhibited a clear mode at much smaller durations (~50-100 ms) with a heavy right handed tail, similar to the histogram reported earlier for achromatic stimuli [15]. Similar results were obtained from M2 (Figure 1D-E).

Figure 2 shows the median burst durations computed for each hue with respect to the gamma power along the horizontal. The median burst duration averaged across all stimuli were clustered around ~300 ms irrespective of the induced gamma power (303.18±23.08 ms for M1 and 312.54±29.50 ms for M2), comparable to the burst durations previously reported for achromatic stimuli [15].

### Gamma bursts in noisy JS model

Next, we tested whether addition of suitable noise in the previously developed JS model ([11,13]; see equation (1)) could reproduce the temporal characteristics of gamma oscillations described above. Presynaptic spikes arriving as inputs to a population are typically modelled as Poisson processes, which have rapid temporal fluctuations. However, dynamics of synaptic and leaky membrane conductances result in smoother temporal profiles. While some previous studies using rate models have used temporally uncorrelated Poisson noise [12,16], we instead used Ornstein-Uhlenbeck (OU) noise [23,24] to approximate the aforementioned factors and to allow independent control over the mean and variance of the noise, which are otherwise coupled for a Poisson process (this is further elaborated in the “*Differential dynamics of input drives to Excitatory and Inhibitory populations”* section in Discussion).

The OU noise formulation (equation (2)) comprises of a temporal summation parameter θ and a pure variance parameter σ of the noise on the oscillations. This noise is described as additive white Gaussian (AWG) noise that is passed through a first order low-pass filter (equation (2)). The parameter θ corresponds to the cutoff frequency of the noisy input (Figure S1). As a result, larger θ results in faster rise in input from baseline (Figure S1A) and permits larger and more rapid (higher frequency) fluctuations in the final noisy input to the corresponding JS population (Figure S1B). OU noise approximates AWG noise as θ approaches infinity. Varying parameter σ, on the other hand, only changes the variance of the final noisy input and has no effect on the transient (Figure S2).

We first studied the effect of varying θ on gamma bursts and transients. Figure 3A shows the time-frequency spectrum of the LFP proxy signal from a single iteration of the JS model with noisy inputs (θ_E_=16 and θ_I_=1). Stimulus presentation was simulated by increasing mean input drive I and variance parameter σ of both populations from 0 to 8 during the stimulus period (0 to 1500 ms), which resulted in gamma oscillations which were bursty, as expected for fluctuating input breaking up the otherwise smooth limit cycle behavior of JS under constant inputs. Detected bursts and their estimated durations are indicated by white horizontal lines. Figure 3B-C shows median burst duration and gamma peak frequency for different values of θ_E_ and θ_I_. Both burst duration and gamma peak-frequency decreased as either θ_E_ or θ_I_ increased. Decrease in burst duration could be explained by the fact that larger θ allows rapid fluctuations in the input to JS which more frequently disrupt its limit cycles, resulting in shorter bursts of continuous oscillations. Figure 3D shows the trial-averaged time-frequency spectra of the LFP proxy signal for different combinations of θ_E_ and θ_I_. In each case, the peak frequency is marked by a white trace over the spectra. Peak frequency reduction (as seen in real LFP data) was replicated when θ_I_ was less than θ_E_. This happens because lower θ_I_ results in slower rise in the input to the I population compared to the E population after stimulus onset. Since increasing inhibitory input drive in the JS model results in decreasing gamma frequency [11], a delayed increase in inhibitory input drive gives rise to a transient decrease in gamma peak frequency.

**Figure 3:**
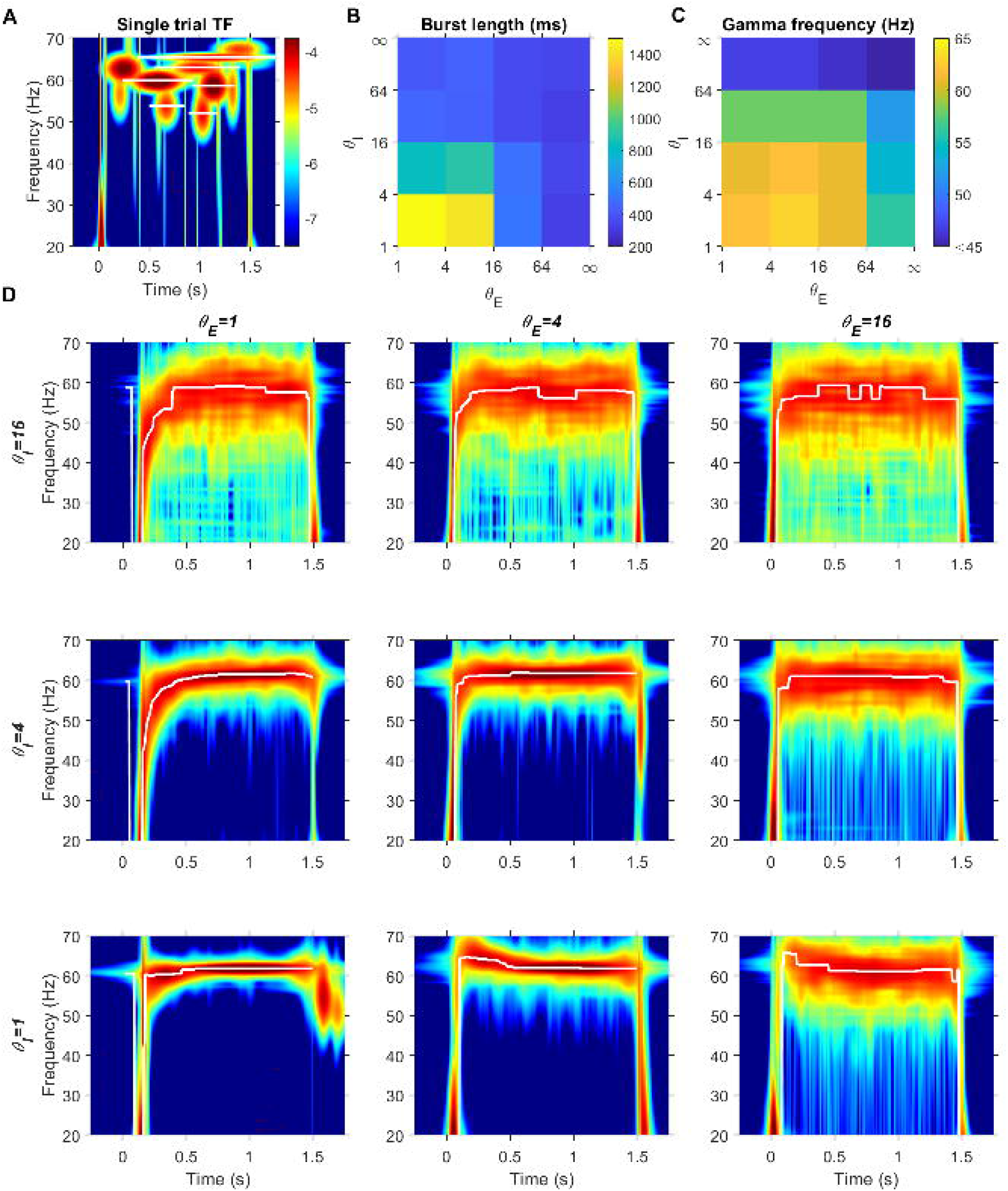
Noisy input parameter θ controls gamma burst duration and peak frequency and determines the slope of the transient. (A) Single trial time-frequency spectrum for a fixed example input combination (I_E_=8, I_I_=8) with θ_E_=16 and θ_I_=1. White horizontal lines indicate the location and durations of detected bursts. (B) Median burst durations across trials for each combination of θ_E_ and θ_I_ for the fixed input combination. θ= ∞ corresponds to the input being ‘Additive White Gaussian’. (C) Gamma peak frequency for each combination of θ_E_ and θ_I_. (D) Trial-averaged time-frequency spectra for different combinations of θ_E_ and θ_I_.

We also studied the effect of varying σ on gamma in the JS with noisy inputs (Figure S3). Both burst duration and gamma peak-frequency decreased as either σ_E_ or σ_I_ increased. Burst duration reduction could be explained by the fact that larger values of σ gave rise to larger fluctuations in the input to JS, which resulted in more disruptions to the default limit cycle behavior of the system and, as a result, shorter burst durations. However, changing σ did not lead to peak frequency reduction after stimulus onset as observed with varying θ. Instead, for higher values of σ_E_ and σ_I_, the gamma power appeared distributed over a wider range of frequencies (larger bandwidth), with a small reduction in the peak frequency (Figure S3D).

### Full model exhibits temporal characteristics of gamma bursts

We chose a model with θ_E_=16 and θ_I_=1, which yielded the desired reduction in peak frequency after stimulus onset as well as the desired burst duration. We constrained the parameter σ of each population to be equal to the square root of the mean input drive (I) to the population (σ_P_^2^=I_P_) to emulate the constant coefficient of variation inherited by these inputs from the Poisson-like presynaptic inputs (i.e., σ_P_^2^ = I_P_ =8 for both populations in equation (2)). Figure 4A shows the distribution of burst durations with a heavy right-sided tail, with median burst duration produced ~250 ms and a mode at ~150 ms. Figure 4B shows a trial-averaged time-frequency spectrum with the peak gamma frequency exhibiting the desired transient decrease.

**Figure 4:**
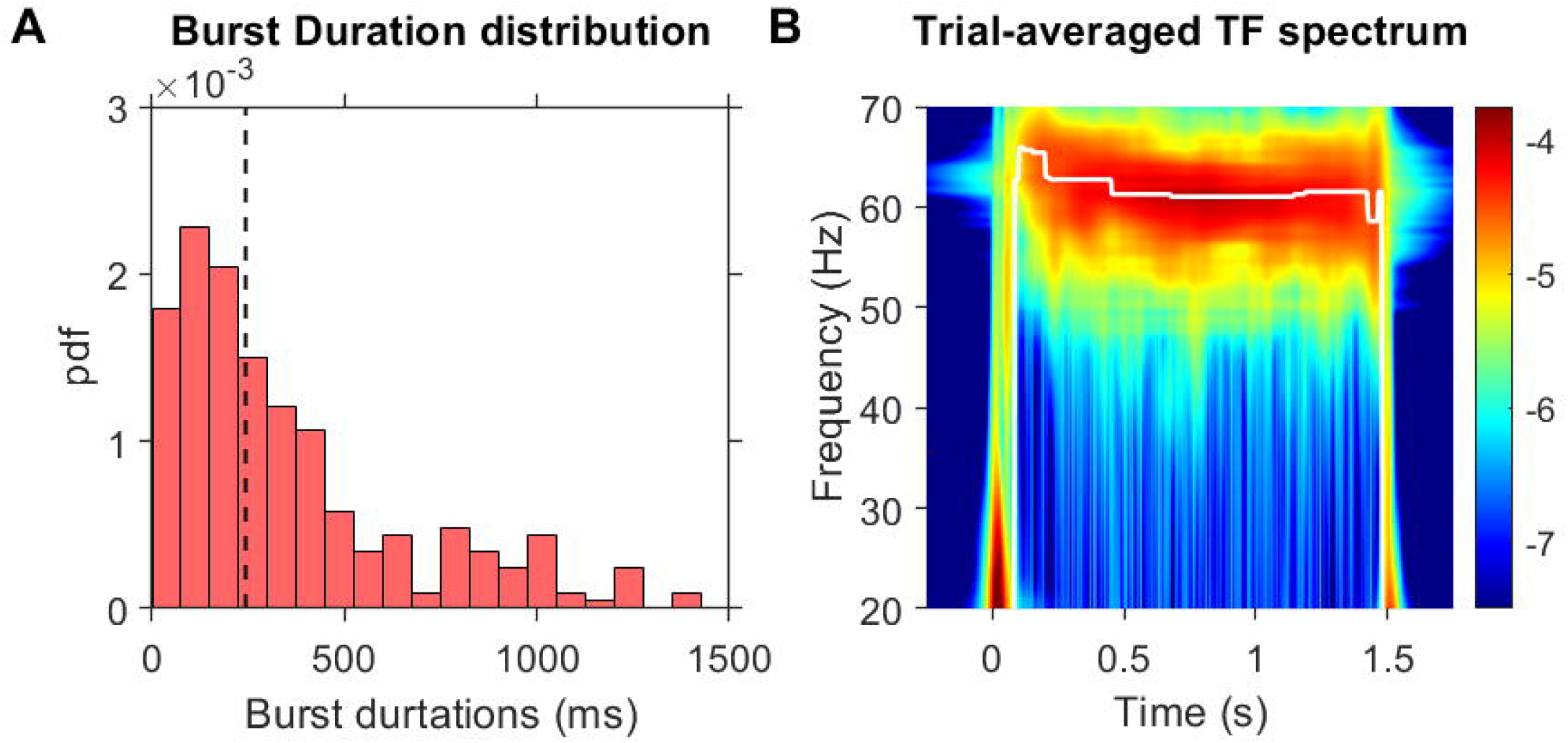
OU noise input with slower onset for inhibitory population demonstrates realistic peak gamma frequency transient and burst length distribution. (A) Probability distribution of gamma burst durations detected in 50 iterations for the condition θ_E_=16 and θ_I_=1 with mean inputs I_E_=8, I_I_=8 and variance parameters σ_E_^2^=8, σ_I_^2^=8. Vertical black line marks the median burst duration detected. (B) Trial-averaged time-frequency spectrum of LFP proxy across 50 iterations with peak gamma frequency.

### Full model exhibits stimulus size and contrast dependence

We tested the behavior of the aforementioned noisy JS model (with θ_E_ and θ_I_ fixed at 16 Hz and 1 Hz) for different input combinations to check the dependence of stimulus size and contrast on gamma (Figure 5). As in previous studies [11,13], presentation of larger size was emulated by increasing inhibitory input drive, which would arise from increased surround activation. Stimuli of higher contrast were emulated by proportionally increasing the input drives to both the populations, corresponding to the increased thalamocortical and lateral inputs to the populations modelled. As before, we set σ_P_^2^ = I_P_ for both populations in equation (3) for each combination since the input combinations scale both the mean and the variance of the effective inputs.

**Figure 5:**
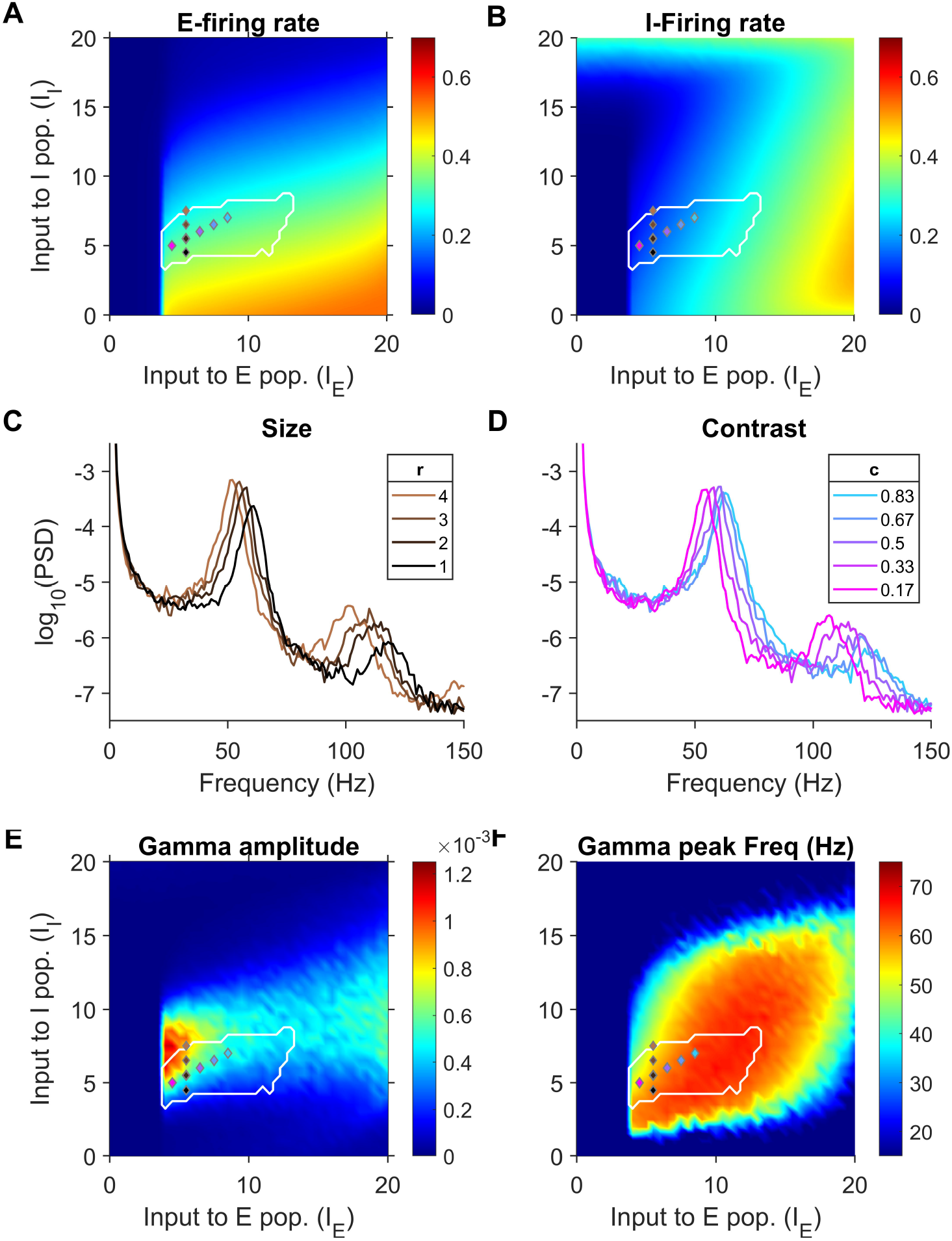
The noisy JS model retains the size-contrast effect of gamma. (A) Mean firing rate of E-population and (B) I-population generated by the JS model with noisy inputs at each input combination. (C) PSD of LFP proxy generated for different stimulus sizes. (D) PSD of LFP proxy generated for different contrast. Input combinations used in (C) and (D) are indicated by markers, with the same colors as traces, in (A), (B) and (E) and (F). (E) Peak gamma amplitude and (F) Peak gamma frequency in the LFP proxy generated for each input combination. The white contour in each figure show the input domain identified by Jadi and Sejnowski (2014) to replicate gamma power increase and peak frequency decrease in response to increasing stimulus size.

Figure 5A-B shows the mean firing rates in the analysis period for different input conditions. The firing rate time series were averaged over 250 to 1250 ms and then this value was averaged across trials and plotted. The white contour encompasses the superlinear regime identified by Jadi and Sejnowski [11] within which size-contrast effects on firing rates and gamma are replicated. Within the white contour regime, increasing size resulted in decrease in firing rates of both populations as desired, while the firing rates increased when excitatory input drive was increased at a faster rate than the inhibitory drive. Hence, for the chosen parameters, varying input drives along such an axis would emulate the contrast effect in firing rates. Figure 5C shows the trial-averaged PSDs for different input combinations within the superlinear regime, emulating increasing sizes by applying larger I_I_. For larger sizes, gamma peak-frequency decreased while gamma power increased, thus replicating the size effect of gamma. Figure 5D shows the PSDs for input combinations within the superlinear regime emulating increasing contrasts. For higher contrasts, gamma frequency increased, as observed in experiments. Figure 5E shows gamma power and Figure 5F shows gamma peak-frequency for all the input combinations. The size-contrast effect on gamma observed in the noiseless case (see Fig 7 of ref [13]) was replicated for the noisy model as well. For Figure 5C-F, the LFP proxy was taken as -r_E_-r_I_ (see ref [13] for more details), but the results remain the same for other proxies as well (data not shown).

### Full model exhibits gamma-harmonic phase difference

Figure 6A shows an example LFP proxy trace computed as -r_E_-r_I_ generated by the full model corresponding to mean input combination of (I_E_=6.5, I_I_=4). The trace shows irregular cycles with sharper troughs resulting in an ‘arch-shape’, as observed in data. In our previous study [13], phase analysis of gamma and its first harmonic in same LFP data revealed a phase difference of 180° between them, and the noiseless JS model could replicate this phase-difference within its JS regime for this LFP proxy. In Figure 6B, the gamma-harmonic phase relationship is presented for the noisy JS model at each input combination with black contours encompassing the ‘arch-shape’ regime within which the input combinations yielded gamma-harmonic phase differences in the range 180±22.5°. The location of the arch-shape regimes were similar to those previously shown for noiseless inputs in each proxy [13], although the extent of these regions were less. In Figure 6C, the mean vector strengths of the phase difference values are shown, with the values ranging between 0.25 to 0.5. These values reveal the consistency of the gamma-harmonic phase difference across subsequent cycles of gamma within each trial. For the JS model with noiseless input, the mean vector strength for all gamma generating inputs was ~1 as the limit-cycle retained a consistent shape in each cycle, giving rise to a fixed gamma-harmonic phase difference at each time-point within a trial (see Fig. S1B of [13]). If the subsequent cycles had less consistent gamma-harmonic phase difference, the vector strength would be lower, with the value of 0 arising when all values of gamma-harmonic phase differences were equally likely to occur at different timepoints within a trial.

**Figure 6:**
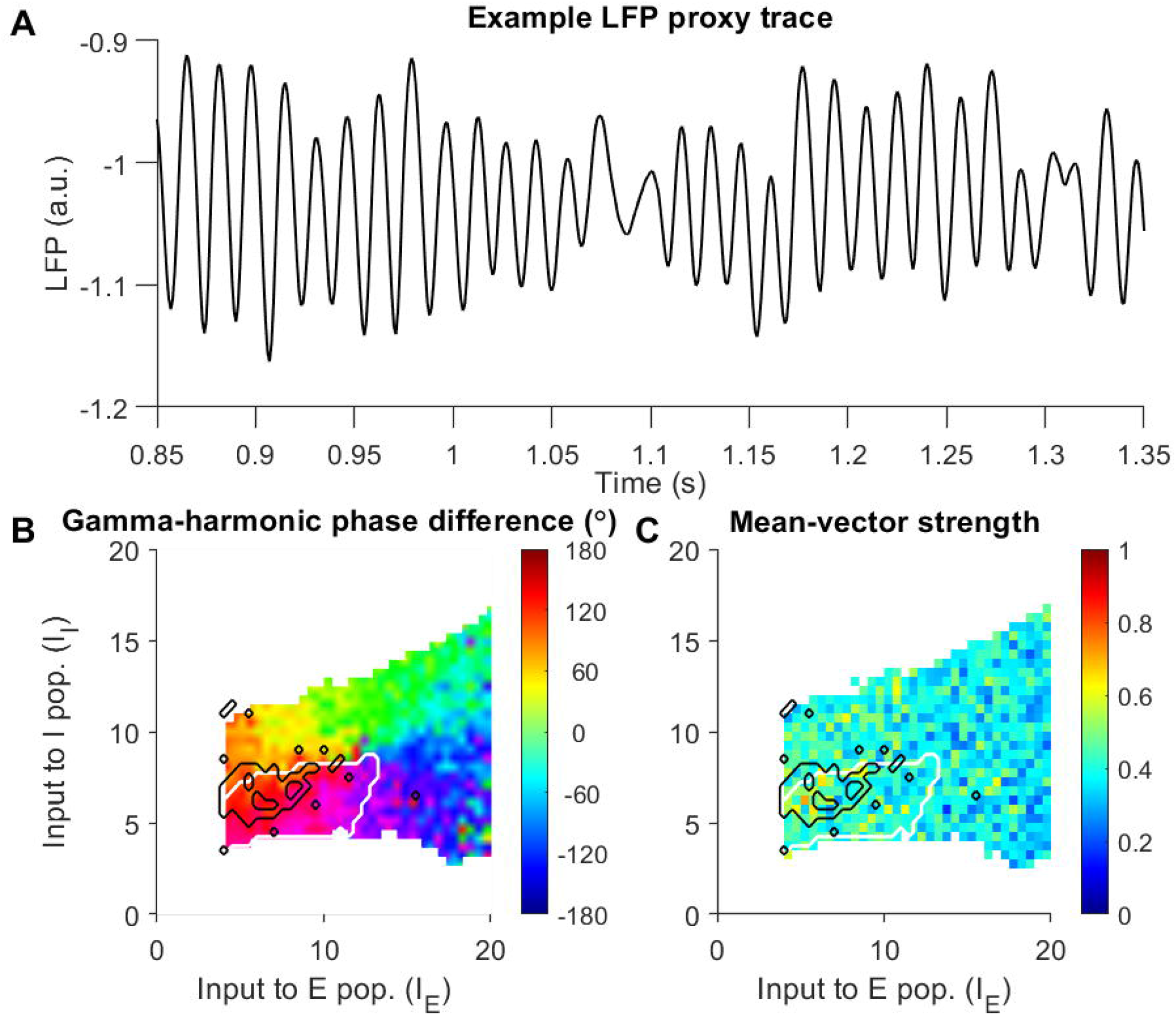
The noisy JS model reproduces the gamma-harmonic phase relationship observed previously in LFP data. (A) Example trace of LFP proxy within the arch-shape regime. (B) Gamma-harmonic phase differences in full noisy JS model for different input combinations. (B) Direction vector strengths of phase differences. Black contours encircle stimulus conditions identified as in the ‘arch-shape regime’. White contours mark the superlinear regime of the JS model.

In summary, the JS model with OU noise successfully demonstrated size-contrast effects and gamma-harmonic phase difference of 180° while also replicating the gamma bursts and transient peak frequency reduction using a suitably slow onset of inhibitory population input drive.

### Poisson noise fails to replicate temporal dynamics and gamma-harmonic phase relationship

We also tested the full model with Poisson inputs. The simulation results for the Poisson inputs over 15 iterations of each input combination are given in Figure S4. In these cases, rhythm burst lengths were very small (Figure S4B), as predicted in Figure 3 for infinite θ, and gamma frequency and power in the model decreased rapidly for higher I_E_ or I_I_. The large variance and zero temporal correlation of this noise gave rise to frequent interruptions to the limit cycle resulting in several short bursts, as evidenced by sharp vertically-oriented gabors in TF spectra, which were centered at a wide range of frequencies. Further, due to the lack of latencies in the input drive, the onset of gamma rhythm was abrupt without any distinguishable slow transient decrease in gamma peak frequency after stimulus onset. The model still exhibited size-effect of gamma as shown in the PSDs plotted in Figure S4C. However, harmonics of gamma were not discernable in the PSDs and the gamma-harmonic phase difference was not consistent within or across trials (data not shown).

## Discussion

We show transient reduction in gamma peak frequency after stimulus onset when different iso-luminant full-screen hues are presented, and show that the burst durations are ~300 ms for different hues, as observed previously for achromatic stimuli [15]. We show that gamma oscillations generated by a self-oscillating WC model of V1 can also show such characteristics when suitable noise is introduced in the input. Further, this noisy JS model also replicates the dependence on stimulus size and contrast, as well as the shape of gamma [13].

### Models of bursty gamma in V1

The bursty nature of gamma could be simulated as noisy limit cycles or quasicycles [23,25]. Noisy limit cycles are produced when the cortical circuitry intrinsically generates sustained oscillations which are modulated by noisy fluctuations. In contrast, quasicycles are obtained when the cortical circuitry dampens eventually to a stable steady state in the absence of further perturbations, exhibiting a spiral trajectory at a specific resonant frequency in gamma range. Quasicycles can be produced by linear models, which are amenable to exact mathematical analyses. Xing and colleagues studied the stochastic nature of V1 gamma in anaesthetized and awake macaques and found a linear noise-driven quasicycle model could be tuned to exhibit the joint distribution of burst frequencies and durations in both conditions [16]. Jia and colleagues expanded this model to include feedback from global excitation which could replicate the stimulus size and contrast dependence on firing rates and gamma rhythm [12]. Linear models lend themselves well to rigorous analyses that can scale well to modelling large populations of several individually defined neurons. For instance, Kang and colleagues [22] constructed a spatial neuronal population model with noisy inputs exhibiting orientation selectivity and resonant bursty gamma showing a reduction in peak frequency with larger stimulus size. Further, a linear auto-regressive model (equivalent to the linear 2D rate models) that was fit to the power spectra of LFP exhibited bursty gamma with realistic relationship between per-cycle amplitude and duration [17]. Similarly, a non-linear damped oscillating network WC model with noisy inputs was also shown to exhibit bursty gamma with the aforementioned burst frequency and duration joint distributions [26]. But oscillations were driven by strong noisy fluctuations preventing the system from converging and thus could not generate prominent harmonics [13,25] as seen in non-sinusoidal gamma found in LFP data. The non-linear JS model used in the current study generates sustained gamma rhythm of constant amplitude, in the absence of noise, through its limit cycles and exhibits appropriate harmonics. The noisy inputs in our current model generated bursts when occasional large fluctuations abruptly change the phase of the rhythm, while the non-linearities of the system largely determine the non-sinusoidal shape and dynamics of the unperturbed cycle period. The noisy JS model, operating as ISN [27], functions analogously to the PING mechanism as both circuits are driven by strong recurrent excitation. Spiking network models [8,28] and stochastic firing network models [25] of PING gamma with Poisson inputs emulating incoming spikes from presynaptic regions have been demonstrated to exhibit bursty gamma rhythms. Wallace and colleagues [25] studied stochastic rate models which approximate well into WC equations in the noisy limit-cycle and quasi-cycle modes and found that while both the models exhibited similar bursty properties, the noisy limit-cycle exhibited a prominent harmonic in the network model and the two-dimensional approximation, which is also observed in our LFP data as shown in our previous study [13]. They also note that while WC approximation to these networks is found to well-approximate uniformly connected networks, those with sparse connectivity introduce additional noise terms in the approximation. This noise is deterministic in origin as it arises from the non-uniformity of connectivity throughout the network and suggests a deterministic chaos component to bursty rhythms in realistic sparse networks, which is not explicitly modelled in the current study. For instance, since a sparsely connected network would contain linked clusters of densely-interconnected sub-networks, each sub-network oscillating at gamma frequency might have slightly different phase from other sub-networks excited by a large stimulus. The sparse lateral interconnections between such sub-networks might result in the lateral input containing a gamma component with slight differences in frequency [29]. However, in our current model, the input drive i_I_ which accounts for lateral input from the surround is asynchronous.

### Differential dynamics of input drives to Excitatory and Inhibitory populations

The desired gamma burst duration and reduction in peak frequency after stimulus onset are both achieved by simply assuming θ_E_ >> θ_I_ in the OU noise model, as the lower value of θ_I_ compared to θ_E_ delays the onset of inhibition, which leads to an initially high peak frequency which reduces as inhibition builds up.

How realistic are these assumptions? In general, excitation and inhibition are tightly balanced, with inhibition generally lagging excitation by a few milliseconds [30,31]. In the classic PING model of gamma generation, incoming excitation excites a local inter-neuronal network (consisting of parvalbumin-positive, soma targeting interneurons), which subsequently shuts down the entire circuitry. This cycle repeats when the inhibition fades away and the E population fires again. The inter-neuronal network is thought to be “local” (acting within a cortical column), although they may send lateral inputs to similarly-tuned surround populations [32]. Generally, gamma oscillations in V1 have been observed to be coherent over 2-3 millimeters (see Fig 8B,D of [33] and Fig 8D of [19]). Therefore, while the incoming excitation to a particular area will depend on the extent of the feedforward projection from the immediately upstream area (LGN in case of V1 neurons), the inhibition will depend on the activation of this larger inhibitory network. This will cause the inhibitory input to be both delayed as well as smoothed. Indeed, stimulus-selective surround suppression is indeed found to have a delayed onset in LFP recordings, emerging ~10 ms after the evoked response onset [34,35]. The lateral connections from surround to the inhibitory neurons have longer ranges [36] and, being unmyelinated, should arrive at different lags depending on the source distance.

This could result in a further delay in the onset of input drive to inhibitory population, as required by our model to exhibit transient decreases in firing rates and gamma frequency after stimulus onset. In our model, the deterministic component of the input drives is a first order linear system. Hence, on an average, input drives evolve gradually and by some time ‘t’ seconds from stimulus onset would have crossed roughly a proportion of (1 − *e*^−(2*πθ*)*t*^) of the range between the initial and steady-state values. Hence, although there is no explicit latency after stimulus onset, the time ‘t’ taken for mean input drive to cross 20% of the range, i.e., (1 − *e*^−(2*πθ*)*t*^ = 0.20), could be used as a proxy. This yields a latency measure given by 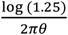 seconds. For the values chosen in the full model (θ_E_=16, θ_I_=1), the half-times of i_E_ and i_I_ are ~2.22 ms and 35.51 ms respectively, indicating a surround suppression latency of ~33 ms.

### Bursty gamma in cortical networks

Some studies have suggested the deterministic source of bursty gamma as a mechanism of visual information processing. Since gamma is strongly modulated by attention [37–40] and its power is found to be decreased with aging and mild cognitive impairments [41], it is hypothesized that gamma entrainment of neuronal firing could be critical for coding of complex stimuli and inter-cortical information transfer [37,42]. Lowet and colleagues found in experiments using gratings of spatially non-uniform contrast that nearby areas oscillating at slightly different frequencies entrain each other through intermittent sharp phase changes which result in interrupted bursts of gamma oscillation and even found similar behavior in a PING model [29]. They further demonstrated through a weakly-coupled oscillator network model that such entrainment is characteristic of such networks. Similarly, Roberts and colleagues had earlier found gamma entrainment across visual areas through intermittent phase shifts followed by concurrent bursts in the local gamma rhythms [6]. Akam and Kullmann have demonstrated selective information routing through entrainment using spiking network models of multiple populations under certain connectivity constraints as a mechanism underlying attention [43].

Our rate model suggests that the bursty nature of gamma could be due to the delayed accumulation of lateral inputs and may not play any functional role. While the potential role of gamma in communication or coding remains debated ({Fries, 2015; Ray and Maunsell, 2015; Ray, 2022}), the temporal dynamics of gamma rhythms provide valuation information about the underlying neural circuitry and properties of neural noise.

## Materials and Methods

### Data Acquisition

LFP data used in this study was recorded by Shirhatti and Ray [20] from V1 of two female macaques, referred to as M1 and M2 in this paper (correspond to M1 and M3 in the earlier paper). As in our recent report [13], LFP from the third monkey was not considered since gamma was relatively weak.

The monkeys performed a passive fixation task as we recorded from a Utah array (96 and 81 electrodes for M1 and M2), which had been implanted in V1 after fitting a titanium headpost on the skull (details of the surgery and implants are provided in [20]), The recording of raw signals from microelectrodes was performed using the 128-channel Cerebus neural signal processor (Blackrock Microsystems). LFP was obtained from the raw signals by filtering online between 0.3 Hz and 500 Hz (Butterworth filters; first-order analog and fourth-order digital respectively) and was recorded at 2 kHz sampling rate and 16-bit resolution and used directly in our analyses.

### Experimental setup and behavior

During the experiment, the subject had its head held still by the headpost as it sat in a monkey chair and viewed a monitor (BenQ XL2411, LCD, 1,280 × 720 resolution, 100 Hz refresh rate) placed ~50 cm from its eyes. A Faraday enclosure housed the subject and the display setup with a dedicated ground for isolation from external electrical noise. The monitor was calibrated and gamma-corrected using i1Display Pro (x-rite PANTONE). Mean luminance was set to 60 cd/m^2^ at the monitor surface and gamma was set to unity for each of the three primaries. The CIE chromaticity xy coordinates of the primaries were: red, (0.644, 0.331); green, (0.327, 0.607); blue, (0.160, 0.062) with the white point at (0.345, 0.358).

The monkeys were subject to a passive fixation task, in which they were rewarded with juice for fixating at a small dot of 0.05°–0.10° radius at the center of the screen throughout the trial (3.3 or 4.8 s duration; fixation spot was displayed throughout). Each trial began with fixation, followed by an initial blank grey screen for a 1,000 ms duration, and then, two to three stimuli were shown for 800 ms each with an interstimulus interval of 700 ms. Juice was awarded only if fixation was maintained within 2° from the fixation point. Trials in which fixation was broken were not considered in our analyses. Eye position data was recorded using the ETL-200 Primate Eye Tracking System (ISCAN) which gave horizontal and vertical coordinates. During the task, monitoring of eye position data, control of task flow, generation and randomization of stimuli were performed using a custom software on macOS. Although fixation was required to be within 2° from the fixation spot, the monkeys were able to hold their fixation well within these limits during the task with a standard deviation of less than 0.4° across all sessions for both monkeys. These small eye movements are unlikely to affect our results since full-screen stimuli were used.

### Stimuli

The set of stimuli comprised of fullscreen patches of 36 hues and 1 achromatic grating. The hues were equally spaced along the circular hue space of the standard HSV nomenclature (0° hue to 350° hue, where 0°, 120°, and 240° correspond to red, green and blue respectively) and were presented at full saturation and value. The achromatic grating was at an orientation of 90° and had a spatial frequency of 4 cpd for M1 and 2 cpd for M2. The grating parameters were chosen to induce strong fast gamma and reduced slow gamma [19]. These full-screen stimuli subtended a visual angle of ~56° in the horizontal direction and ~33° in the vertical direction.

### Electrode selection

As with our previous reports [13,20], we considered only those electrodes for analysis that gave consistent stimulus-induced changes and reliable receptive field estimates across sessions. These criteria were determined by a receptive field mapping protocol that was run across multiple days [44]. Furthermore, electrodes with unusable or inconsistent signals, a high degree of crosstalk with other electrodes, or impedances outside the range of 250–2,500 KΩ for monkey M1 and 125–2,500 KΩ for M2 were discarded, finally yielding 64 and 16 electrodes from M1 and M2, respectively.

### Data Analysis

We discarded stimulus presentations with excessive artifacts for each session (<5.9% and <5.0% of presentations of each stimulus in M1 and M2), yielding 19.9±6.2 trials in M1 and 20.2±1.0 trials in M2 per stimulus as in our previous studies [13,20].

### Time-Frequency analysis of LFP

Time-Frequency spectrum was computed using Matching Pursuit (MP) [45]. The raw LFP signal was filtered through a Butterworth filter (zero-phase; order 4) with a passband of 20-80 Hz. The filtered signal was subject to MP analysis with a stochastic dictionary, which iteratively fits gabors to the LFP traces and computes the time frequency spectrum of the traces by adding together the time-frequency spectra of all these gabors (see [15] for details). Prior to MP, Low pass filtering was done to reduce computational load during MP; TF results within the gamma band were similar for higher passbands. In each trial, for a given electrode, gabors which were centered within the stimulus presentation duration (0 to 800 ms from stimulus onset) and had their center frequencies within 20-80 Hz were considered as ‘gamma bursts’ and burst durations were computed from the scales of the respective gabors (refer [15] for more details). Further, bursts which started before 0 ms were discarded.

To compute the electrode averaged time-frequency power spectrum, for each stimulus, the time-frequency spectrum from each trial corresponding to the presentation of the stimulus from each electrode were averaged. Then the baseline power at each frequency was computed by averaging the power at the respective frequency over the baseline period interval between - 100 ms to 0 ms from stimulus onset. This baseline for each frequency was subtracted (on a log scale) from the averaged time-frequency spectrum at every timepoint at the frequency. The peak gamma frequency time series was computed for each stimulus as the frequency within 20 to 80 Hz range with the maximum power in the electrode averaged time frequency spectrum at each timepoint.

### Spectral analysis of LFP

For computing the electrode averaged Power Spectral Density (PSD) for each stimulus, the LFP recorded from −500 to 0 ms from stimulus onset was taken as the ‘baseline period’ and 250 to 750 ms from stimulus onset was taken as the ‘stimulus period’ to avoid the transient responses to the stimulus onset. This yielded a frequency resolution of 2 Hz in the PSD which was computed using the Multitaper method using the Chronux toolbox [46], with three tapers. The change in power was calculated as 10 times the difference between base-10 logarithm of PSDs at stimulus period and baseline period, expressed in decibels (dB). Estimation of peak frequencies was done on these baseline-corrected PSDs.

Gamma range was taken as 20-80 Hz, and ‘gamma peak frequency’ was estimated as the highest peak within this range and the height of the peak was taken as gamma power for the corresponding stimulus.

### Experimental Design and Statistical Analysis

To estimate gamma burst durations, detected bursts from each trial of the chosen stimulus presentation were pooled across all electrodes and the median of the resulting distribution was computed and plotted in Figure 2. To compute an errorbar to the median for each stimulus, a standard error was estimated by bootstrapping over N iterations (where N is the number of datapoints). This involved random sampling with replacement of the ratio data N times and estimating their median each time, which resulted in N medians, whose standard deviation is taken as the length of each half of the errorbar. The standard error (SE) was taken as the standard deviation of the median burst durations from all the stimuli.

### Jadi-Sejnowski (JS) model

Jadi and Sejnowski [11] used a simple rate model consisting of an excitatory and an inhibitory population with sigmoidal activation, operating as an Inhibition Stabilized Network (ISN) and constrained the input drives to the populations to reproduce the increase in power and decrease in peak frequency of gamma with increasing stimulus size as earlier studies have observed in V1 [2,3].

The model defines the population firing rates r_E_ and r_I_ of Excitatory and Inhibitory populations respectively as follows:

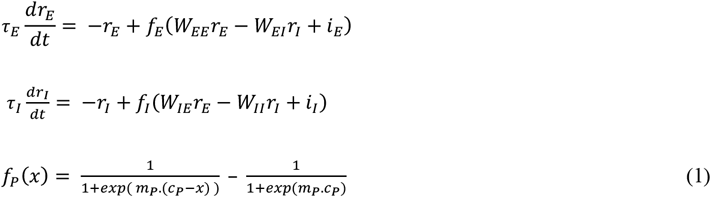

The JS model originally used constant input drives i_E_ and i_I_. In this model, larger stimuli cause an increased inhibitory drive (i_I_), owing to suppression from the surrounding populations. Since the model operated close to a supercritical Hopf bifurcation, the amplitude and frequency of oscillations in firing rates could be closely approximated by linearization of the model. The authors deduced and demonstrated that the model gave rise to the observed trends in the gamma when the inputs were such that the inhibitory population was strongly ‘superlinear’. This means that the summed inputs to the inhibitory population (from recurrent and external sources; argument of f_I_ in equation (1)) must lie in a certain range of values where the activation function σ_I_ curves upwards (increasing in slope with increasing summed input). For the sigmoidal activation function used in equation (1), the summed inputs must operate in the lower half of the sigmoid. Superlinear activation of the excitatory population, on the other hand, was antagonistic (not strongly superlinear). The set of such inputs constitute the operating regime of the model, which we refer to as the ‘superlinear’ regime. The parameter combination used in this study is given in Table 1. The connectivity weights were retained as that of the original model by Jadi and Sejnowski [11] but c_E_ and c_I_ (θ_E_ and θ_I_ in [11]) were changed to ensure that our baseline period input combination (i_E_,i_I_) = (0,0) does not have multiple fixed points but yields steady-state value of firing rates close to 0. Further, the time-constants τ_E_ and τ_I_ were scaled by a factor of 10/13=0.769 from the original parameters to ensure that the model generates a wide range of limit cycles frequencies spanning the gamma range.

**Table 1:**
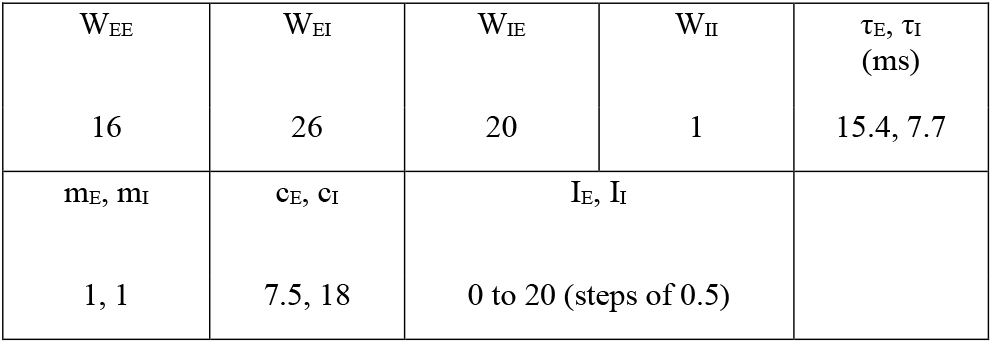
Parameter values used in JS model with noisy inputs.

In our study, we subject the model to noisy input drives i_E_ and i_I_, which were both Ornstein-Uhlenbeck (OU) noise. The OU noise formulation of i_E_ and i_I_ was a first-order low pass filtered white Gaussian noise:

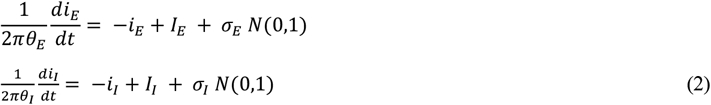

where I_E_ and I_I_ denote the desired steady-state mean values of i_E_ and i_I_ respectively, and *N*(0,1) denotes that at each timepoint the value is sampled from a Normal distribution.

The above formulation could be considered as an Additive White Gaussian (AWG) noise passed through a first-order low pass filter with a cutoff frequency at θ_P_ for a population P. Further, at steady state, the variance would be *πσ*_*P*_^2^*θ*_*P*_.

As θ_P_ is increased to infinity, the input drive tends to AWG noise and the formulation is as follows:

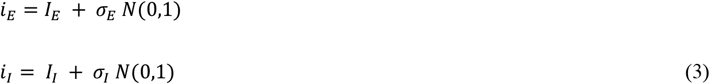

Finally, we simulated a ‘full noise-driven JS model’ to demonstrate bursty gamma and replicate the size-contrast effects on gamma using OU inputs as in equation (2), constrained such that the variance equals the mean of the input gaussian on the right-hand side of the equation (3), i.e., σ_P_^2^ = I_P_. This would ensure that the steady-state variance of i and i would be proportional to their steady-state mean I_E_ and I_I_ respectively.

We also tested the full JS model with Poisson inputs with means I_E_ and I_I_ respectively (Figure S4). The inputs were formulated as follows:

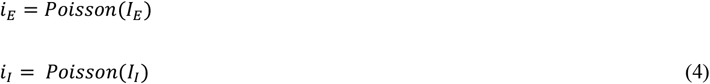

### Spectral analysis of model output

Since the model was used with stochastic inputs, each input combination was simulated for 50 repeats. Each repeat spanned from −2.047 to 2.048 s and was simulated by the forward Euler method with a step size of 2e-5 s. The steady-state mean inputs I_E_ and I_I_ and variances σ_E_ and σ_I_ were maintained at 0 from −2.047 to 0 s, then instantly the test inputs and variances were applied from 0 to 1.5 s, followed by instantly returning inputs and variances to 0. This simulates a stimulus presentation for 1.5 s. The model firing rate time series was then passed through a low pass Butterworth filter (order 4) with cutoff at 200 Hz and downsampled by 50 to a sampling frequency of 1 kHz with 4096 timepoints in each trial. In further analyses, ‘stimulus period’ was taken to be 0.25 to 1.25 ms to avoid transients from input onset and offset. An LFP proxy was computed from r_E_ and r_I_. LFP proxies considered were -r_E_-r_I_, -r_E_ and -r_I_ (as in our previous study [13]).

Time frequency spectrum of the LFP proxy was computed using the Matching Pursuit algorithm (MP) with a stochastic dictionary in the same way as for the data, taking the stimulus period to be from 250 to 1250 ms and the baseline from −250 to 0 ms. Gabors detected within the TF stimulus period were taken as ‘bursts’ and the duration was computed from the scale of the gabor. The burst duration data presented in the results were from the LFP proxy -r_E_-r_I_. However, the results were similar for other proxies. Further, trial-averaged PSD was computed for each stimulus and parameter combination by the Multitaper method using the Chronux toolbox [46], over the interval 250 to 1250 ms. The maximum power in the PSD in the range 30 to 70 Hz was taken as the gamma peak and the corresponding frequency as the peak frequency.

### Harmonic analysis of model output

To compute the phase difference of gamma and its first harmonic in the simulation output, the trial-averaged PSD of an LFP proxy was computed for each stimulus combination Gamma and harmonic signals were extracted from the LFP proxy signal during the stimulus period by bandpass filtering using separate Butterworth filters (zero-phase; order 4). The passband for gamma was 20 Hz wide, centered around the gamma peak frequency, identified from trial-averaged PSD for the input combination. The passband for the first harmonic of gamma was also 20 Hz wide but centered around twice the corresponding gamma peak frequency. The phases of these signals (φ_gamma_ and φ_harmonic_) were then computed using Hilbert transform. The phase difference between gamma and its harmonic was calculated as:

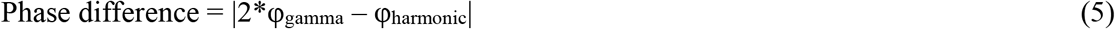

The above phase difference estimate was computed at each timepoint within the stimulus period of every trial. Note that the 20 Hz passband for gamma and its harmonic-band filters is wide enough so that errors in gamma peak-frequency detection do not affect the phase difference estimates. Circular statistics on the gamma-harmonic phase difference data were computed in Matlab using the Circular Statistics toolbox [47]. The mean phase difference is the circular mean (implemented in circ_mean function of the Circular Statistics toolbox).

To validate that the distribution of gamma-harmonic phase differences was non-uniform and, thus, assess the validity of circular mean estimates in Figure 6, we subjected phase-difference from each trial for a given input to a Rayleigh test of non-uniformity (Null hypothesis: uniform distribution of phases). When the gamma activity recorded for a given input has a specific waveform, the corresponding distribution of gamma-harmonic phase-differences will be unimodal. We considered this to be the case when the p-value of the Rayleigh test was less than 0.01. Among the inputs with unimodal phase-differences, those with average phase-difference of 180±22.5° across trials were considered to be constitute the ‘arch-shape regime’ in Figure 6 as the phase-difference in recorded LFP data is found to be around 180° (refer [13]), Furthermore, we computed the mean vector strength of the gamma-harmonic phase-difference to quantify the consistency of the non-sinusoidal distortion in subsequent gamma cycles. At each time point within the analysis window in a trial, unit vectors were taken with orientation equal to the gamma-harmonic phase-difference at each time point within the analysis window and the magnitude of the average of these unit vectors was considered as the mean vector strength for the corresponding trial. The mean vector strengths across trials of a specific input combination were then averaged.

## Figures

**Figure S1:**
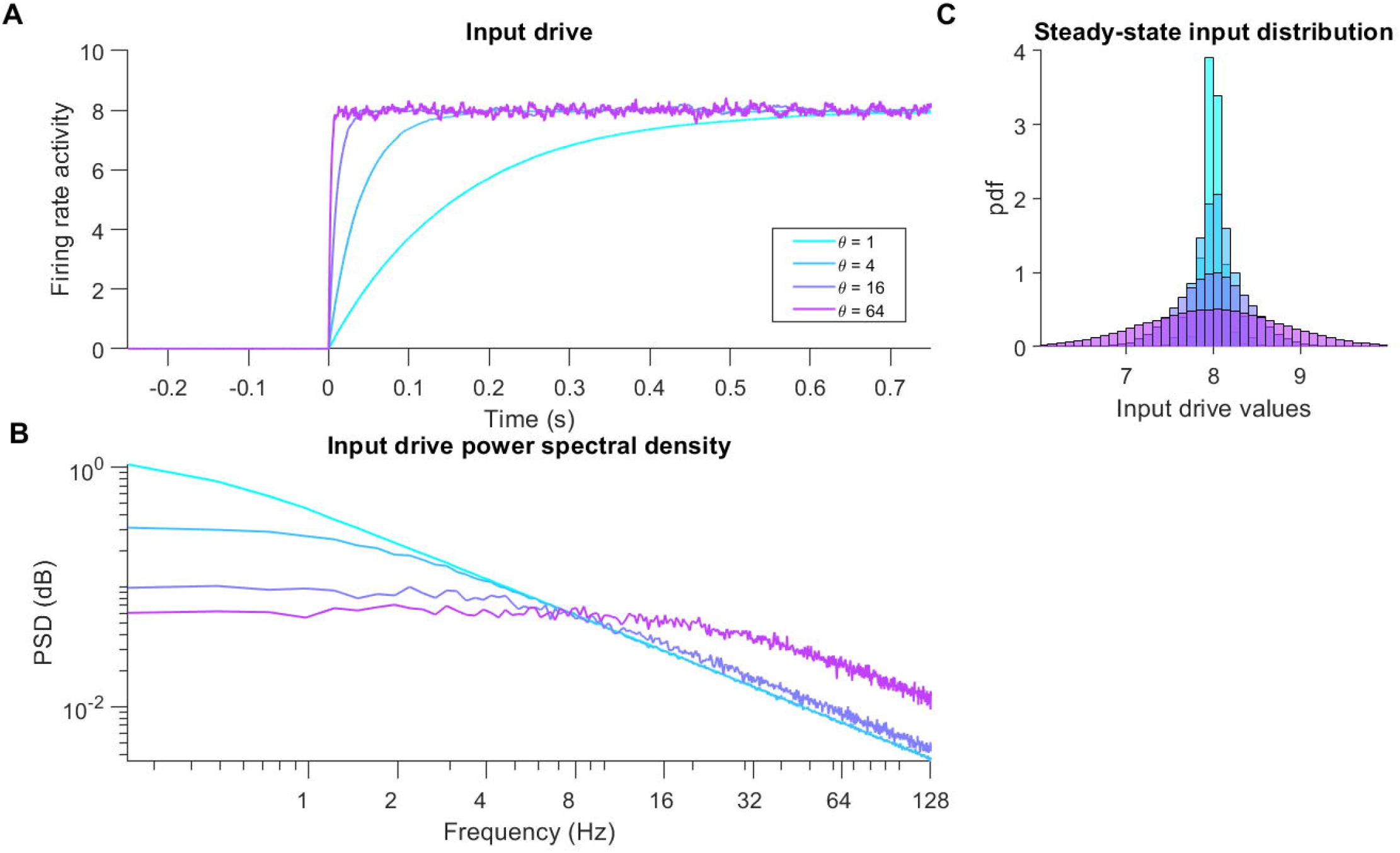
Increasing θ increases the cutoff frequency input drive resulting in faster transients and higher fluctuations in the input drive. (A) Mean input noise time-series computed from 50 OU noise input drive signals generated for each value of θ. (B) Distributions of input signal values within the 1-1.5 s interval across 50 iterations for each θ. (C) Average power spectral density across 50 iterations of the OU noise input signal.

**Figure S2:**
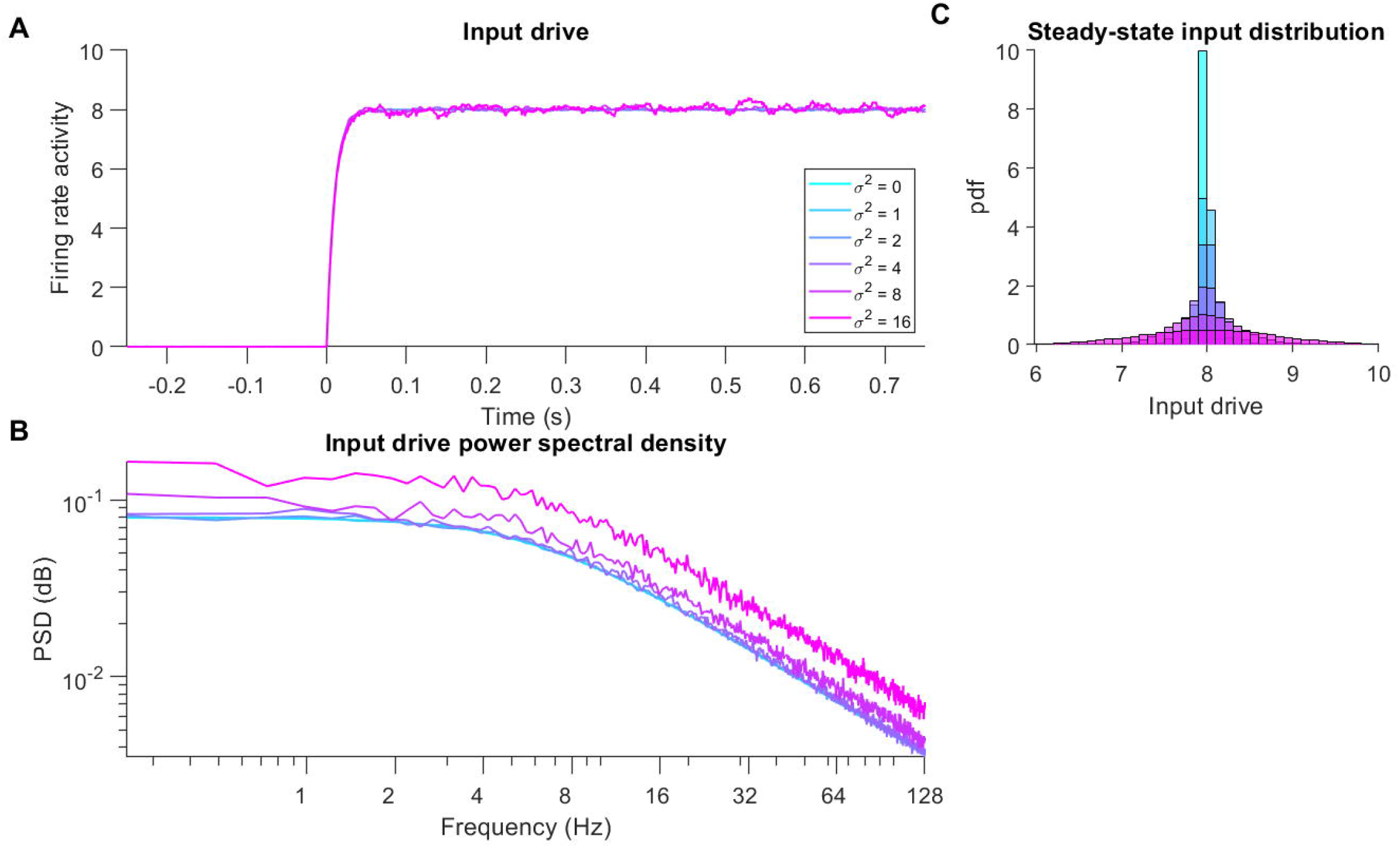
Increasing σ scales up noisy fluctuations in the input drive without affecting the temporal characteristics. (A) Mean input noise time-series computed from 50 OU noise input drive signals generated for each value of σ. (B) Distributions of input signal values within the 1-1.5 s interval across 50 iterations for each σ. (C) Average power spectral density across 50 iterations of the OU noise input signal.

**Figure S3:**
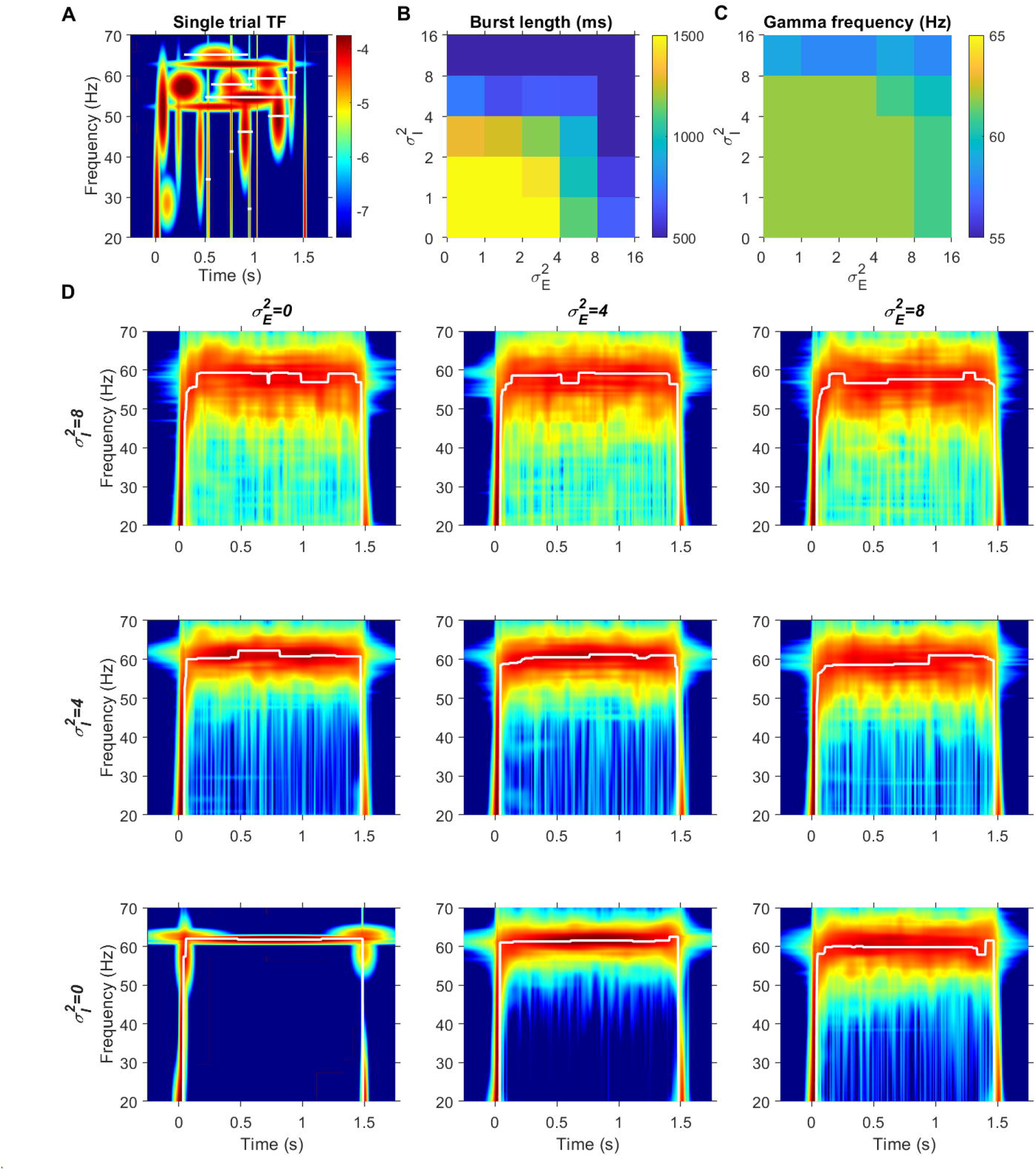
Noisy input parameter σ decreases gamma burst duration and peak frequency but does not determine the slope of the transient. (A) Single trial time-frequency spectrum for a fixed example input combination (I_E_=8, I_I_=8) and fixed θ’s (θ_E_=16, θ_I_=16) with σ_E_^2^=8 and σ_I_^2^=8. White horizontal lines indicate the location and durations of detected bursts. (B) Median burst durations across trials for each combination of θ_E_ and θ_I_ for the fixed example input combination (I_E_=8, I_I_=8) and fixed θ’s (θ_E_=16, θ_I_=1). (C) Gamma peak frequency for each combination of σ_E_ and σ_I_. (D) Trial-averaged time-frequency spectra with peak frequency of gamma marked by white trace for different combinations of σ_E_ and σ_I_.

**Figure S4:**
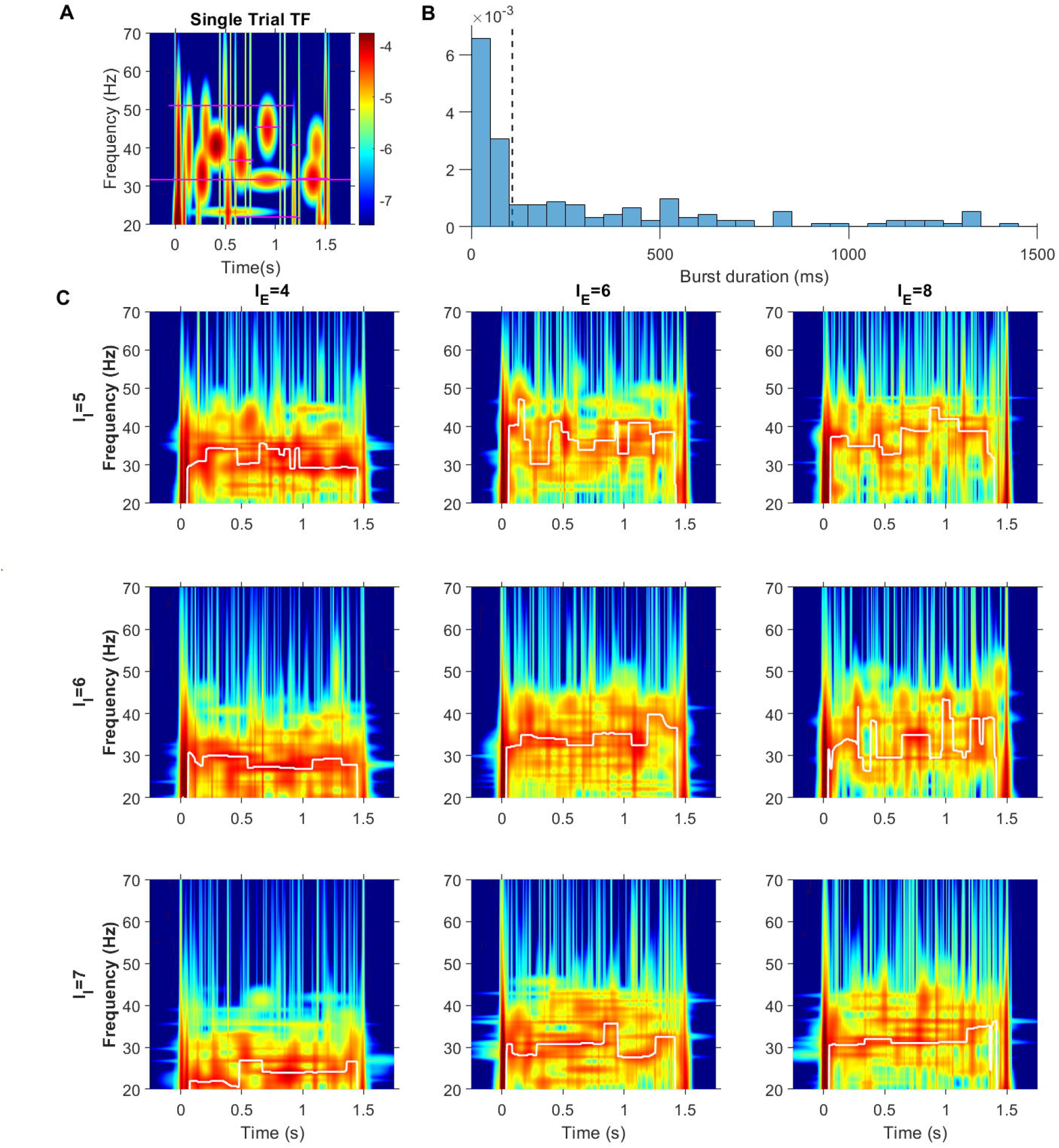
Poisson inputs induce very short bursts in the JS model due to their high variance and rapid fluctuations with zero temporal correlation. (A) Time frequency power spectrum of LFP proxy from single trial with I_E_=6 and I_I_=7. Detected bursts are marked by horizontal magenta lines whose length indicates the estimated burst duration. (B) Probability distribution of burst durations across 15 iterations of the example input combination. (C) Trial-averaged time-frequency spectra of LFP proxy for different combinations of I_E_ and I_I_. White traces mark the peak gamma frequency at each timepoint.

## References

1. Wang XJ. Neurophysiological and Computational Principles of Cortical Rhythms in Cognition. Physiol Rev. 2010 Jul 1;90(3):1195–268.

2. Gieselmann MA, Thiele A. Comparison of spatial integration and surround suppression characteristics in spiking activity and the local field potential in macaque V1. Eur J Neurosci. 2008 Aug;28(3):447–59.

3. Ray S, Maunsell JH. Different origins of gamma rhythm and high-gamma activity in macaque visual cortex. PLoS Biol. 2011;9(4).

4. Peter A, Uran C, Klon-Lipok J, Roese R, van Stijn S, Barnes W, et al. Surface color and predictability determine contextual modulation of V1 firing and gamma oscillations. Colgin L, editor. eLife. 2019 Feb 4;8:e42101.

5. Ray S, Maunsell JHR. Differences in Gamma Frequencies across Visual Cortex Restrict Their Possible Use in Computation. Neuron. 2010 Sep 9;67(5):885–96.

6. Roberts MJ, Lowet E, Brunet NM, Ter Wal M, Tiesinga P, Fries P, et al. Robust Gamma Coherence between Macaque V1 and V2 by Dynamic Frequency Matching. Neuron. 2013 May 8;78(3):523–36.

7. Buzsáki G, Wang XJ. Mechanisms of Gamma Oscillations. Annu Rev Neurosci. 2012;35(1):203–25.

8. Chariker L, Shapley R, Young LS. Rhythm and Synchrony in a Cortical Network Model. J Neurosci. 2018 Oct 3;38(40):8621–34.

9. Zachariou M, Roberts MJ, Lowet E, De Weerd P, Hadjipapas A. Empirically constrained network models for contrast-dependent modulation of gamma rhythm in V1. NeuroImage. 2021 Apr 1;229:117748.

10. Wilson HR, Cowan JD. Excitatory and inhibitory interactions in localized populations of model neurons. Biophys J. 1972 Jan;12(1):1–24.

11. Jadi MP, Sejnowski TJ. Regulating Cortical Oscillations in an Inhibition-Stabilized Network. Proc IEEE Inst Electr Electron Eng. 2014 Apr 21;102(5).

12. Jia X, Xing D, Kohn A. No consistent relationship between gamma power and peak frequency in macaque primary visual cortex. J Neurosci Off J Soc Neurosci. 2013 Jan 2;33(1):17–25.

13. Krishnakumaran R, Raees M, Ray S. Shape analysis of gamma rhythm supports a superlinear inhibitory regime in an inhibition-stabilized network. PLOS Comput Biol. 2022 Feb 14;18(2):e1009886.

14. Shirhatti V, Ravishankar P, Ray S. Gamma oscillations in primate primary visual cortex are severely attenuated by small stimulus discontinuities. PLOS Biol. 2022 Jun 14;20(6):e3001666.

15. Chandran KS S, Seelamantula CS, Ray S. Duration analysis using matching pursuit algorithm reveals longer bouts of gamma rhythm. J Neurophysiol. 2017 Nov 8;119(3):808–21.

16. Xing D, Shen Y, Burns S, Yeh CI, Shapley R, Li W. Stochastic Generation of Gamma-Band Activity in Primary Visual Cortex of Awake and Anesthetized Monkeys. J Neurosci. 2012 Oct 3;32(40):13873–13880a.

17. Spyropoulos G, Saponati M, Dowdall JR, Schölvinck ML, Bosman CA, Lima B, et al. Spontaneous variability in gamma dynamics described by a damped harmonic oscillator driven by noise. Nat Commun. 2022 Apr 19;13(1):2019.

18. Brunet N, Bosman CA, Roberts M, Oostenveld R, Womelsdorf T, De Weerd P, et al. Visual cortical gamma-band activity during free viewing of natural images. Cereb Cortex N Y N 1991. 2015 Apr;25(4):918–26.

19. Murty DVPS, Shirhatti V, Ravishankar P, Ray S. Large Visual Stimuli Induce Two Distinct Gamma Oscillations in Primate Visual Cortex. J Neurosci Off J Soc Neurosci. 2018 14;38(11):2730–44.

20. Shirhatti V, Ray S. Long-wavelength (reddish) hues induce unusually large gamma oscillations in the primate primary visual cortex. Proc Natl Acad Sci. 2018 Apr 24;115(17):4489–94.

21. Henrie JA, Shapley R. LFP Power Spectra in V1 Cortex: The Graded Effect of Stimulus Contrast. J Neurophysiol. 2005 Jul 1;94(1):479–90.

22. Kang K, Shelley M, Henrie JA, Shapley R. LFP spectral peaks in V1 cortex: network resonance and cortico-cortical feedback. J Comput Neurosci. :13.

23. Powanwe AS, Longtin A. Amplitude-phase description of stochastic neural oscillators across the Hopf bifurcation. Phys Rev Res. 2021 Jul 9;3(3):033040.

24. tuckwell HC, Wan FYM, Rospars JP. A spatial stochastic neuronal model with Ornstein–Uhlenbeck input current. Biol Cybern. 2002 Feb 1;86(2):137–45.

25. Wallace E, Benayoun M, Drongelen W van, Cowan JD. Emergent Oscillations in Networks of Stochastic Spiking Neurons. PLOS ONE. 2011 May 6;6(5):e14804.

26. Powanwe AS, Longtin A. Determinants of Brain Rhythm Burst Statistics. Sci Rep. 2019 Dec 4;9(1):18335.

27. sodyks MV, Skaggs WE, Sejnowski TJ, McNaughton BL. Paradoxical effects of external modulation of inhibitory interneurons. J Neurosci Off J Soc Neurosci. 1997 Jun 1;17(11):4382–8.

28. Mazzoni A, Lindén H, Cuntz H, Lansner A, Panzeri S, Einevoll GT. Computing the Local Field Potential (LFP) from Integrate-and-Fire Network Models. PLOS Comput Biol. 2015 Dec 14;11(12):e1004584.

29. Lowet E, Roberts MJ, Peter A, Gips B, De Weerd P. A quantitative theory of gamma synchronization in macaque V1. Schroeder CE, editor. eLife. 2017 Aug 31;6:e26642.

30. Okun M, Lampl I. Instantaneous correlation of excitation and inhibition during ongoing and sensory-evoked activities. Nat Neurosci. 2008 May;11(5):535–7.

31. Wehr M, Zador AM. Balanced inhibition underlies tuning and sharpens spike timing in auditory cortex. Nature. 2003 Nov;426(6965):442–6.

32. Shushruth S, Mangapathy P, Ichida JM, Bressloff PC, Schwabe L, Angelucci A. Strong Recurrent Networks Compute the Orientation Tuning of Surround Modulation in the Primate Primary Visual Cortex. J Neurosci. 2012 Jan 4;32(1):308–21.

33. Jia X, Smith MA, Kohn A. Stimulus Selectivity and Spatial Coherence of Gamma Components of the Local Field Potential. J Neurosci. 2011 Jun 22;31(25):9390–403.

34. Angelucci A, Bijanzadeh M, Nurminen L, Federer F, Merlin S, Bressloff PC. Circuits and Mechanisms for Surround Modulation in Visual Cortex. Annu Rev Neurosci. 2017;40(1):425–51.

35. Wang T, Li Y, Yang G, Dai W, Yang Y, Han C, et al. Laminar Subnetworks of Response Suppression in Macaque Primary Visual Cortex. J Neurosci. 2020 Sep 23;40(39):7436–50.

36. Girard P, Hupé JM, Bullier J. Feedforward and Feedback Connections Between Areas V1 and V2 of the Monkey Have Similar Rapid Conduction Velocities. J Neurophysiol. 2001 Mar 1;85(3):1328–31.

37. Chalk M, Herrero JL, Gieselmann MA, Delicato LS, Gotthardt S, Thiele A. Attention reduces stimulus-driven gamma frequency oscillations and spike field coherence in V1. Neuron. 2010 Apr 15;66(1):114–25.

38. Bosman CA, Schoffelen JM, Brunet N, Oostenveld R, Bastos AM, Womelsdorf T, et al. Attentional Stimulus Selection through Selective Synchronization between Monkey Visual Areas. Neuron. 2012 Sep 6;75(5):875–88.

39. Das A, Ray S. Effect of Stimulus Contrast and Visual Attention on Spike-Gamma Phase Relationship in Macaque Primary Visual Cortex. Front Comput Neurosci. 2018;12:66.

40. Ferro D, van Kempen J, Boyd M, Panzeri S, Thiele A. Directed information exchange between cortical layers in macaque V1 and V4 and its modulation by selective attention. Proc Natl Acad Sci. 2021 Mar 23;118(12):e2022097118.

41. Murty DVPS, Manikandan K, Kumar WS, Ramesh RG, Purokayastha S, Javali M, et al. Gamma oscillations weaken with age in healthy elderly in human EEG. NeuroImage. 2020 Jul 15;215:116826.

42. Fries P. Rhythms for Cognition: Communication through Coherence. Neuron. 2015 Oct 7;88(1):220–35.

43. Akam T, Kullmann D. Efficient “communication through coherence” requires oscillations structured to minimize interference between signals. PLoS Comput Biol [Internet]. 2012 [cited 2022 Oct 27];8(11). Available from: https://ora.ox.ac.uk/objects/uuid:1e5e3117-7823-42e9-988c-692b2fb6adfb

44. Dubey A, Ray S. Cortical Electrocorticogram (ECoG) Is a Local Signal. J Neurosci. 2019 May 29;39(22):4299–311.

45. Mallat SG, Zhifeng Zhang. Matching pursuits with time-frequency dictionaries. IEEE Trans Signal Process. 1993 Dec;41(12):3397–415.

46. Bokil H, Andrews P, Kulkarni JE, Mehta S, Mitra PP. Chronux: A platform for analyzing neural signals. J Neurosci Methods. 2010 Sep 30;192(1):146–51.

47. Berens P. CircStat: A MATLAB Toolbox for Circular Statistics. J Stat Softw. 2009 Sep 23;31(1):1–21.

